# Quantitative analysis of spore shapes improves identification of fungi

**DOI:** 10.1101/2021.04.08.438929

**Authors:** Alexander Ordynets, Sarah Keßler, Ewald Langer

**Affiliations:** Department of Ecology, Faculty of Mathematics and Natural Sciences, University of Kassel, Germany

**Keywords:** geometric morphometrics, classification, contour, fungi, outline, traditional morphometrics

## Abstract

Morphology of organisms is an important source of evidence for biodiversity assessment, taxonomic decisions, and understanding of evolution. Shape information about zoological and botanical objects is often treated quantitatively and in this form improves species identification. In studies of fungi, quantitative shape analysis was almost ignored. The disseminated propagules of fungi, the spores, are crucial for their taxonomy – currently in the form of linear measurements or subjectively defined shape categories. It remains unclear how much quantifying spore shape information can improve species identification. In this study, we tested the hypothesis that shape, as a richer source of information, overperforms size when performing automated identification of fungal species. We used the fungi of the genus *Subulicystidium* (Agaricomycetes, Basidiomycota) as a study object. We analysed 2D spore shape data via elliptic Fourier and Principal Component analyses. With flexible discriminant analysis, we achieved a slightly higher species identification success rate for shape predictors (61.5%) than for size predictors (59.1%). However, we achieved the highest rate for a combination of both (64.7%). We conclude that quantifying fungal spore shapes is worth the effort. We provide an open access protocol which, we hope, will stimulate a broader use of quantitative shape analysis in fungal taxonomy. We also discuss the challenges of such analyses that are specific to fungal spores.

## Introduction

In eucaryotic organisms, morphology is an important source of evidence for biodiversity assessment, taxonomic decisions, and understanding of evolutionary and ecological processes. Morphological information is quickly accessible and can be processed at relatively low costs. Therefore it is broadly used by both researchers and citizen scientists. Morphological information, on the one hand, provides a starting point for molecular analyses, and on the other hand, serves as reference data to validate the molecular results [1].

Morphology covers two principal concepts: size and shape. The former is easier to measure and was dominating for more than a century in the quantitative analyses known as traditional morphometrics [2]. Although being important, the size alone is often insufficient for the delimitation of species or populations. For example, within a single taxonomic group of diatom algae (Bacillariophyceae), linear measurements allowed to delimit species in some genera [3] but not in others [4]. The authors of the latter study concluded that involving quantitative shape descriptors, in addition to size, would make delimitation of taxa more efficient. It is a discipline of geometric morphometrics that aims to quantify a shape, geometric information about the object that remains after removing the effects of location, rotation, and scale [5]. Tools of geometric morphometrics allow splitting the shape information into symmetric and asymmetric components and analyse them separately [6]. Besides dealing with richer data of numeric nature, geometric morphometric allows reconstruction of the original look of the object after analyses – a valuable property that traditional morphometrics does not offer [1,2].

Fungi are among the species-richest organism groups on Earth [7]. Generally, for morphology-based fungal taxonomy, the features of the disseminated propagules, the spores, are of the highest priority among phenotypic characters [8]. However, there is a difference in how the size and shape of spores are usually treated. The spore size is routinely used for species delimitations, mostly in the form of quantitative traits such as length and width [9,10]. Spore shape also plays a big role but has been treated differently. The length to width ratio is used frequently as a proxy of the spore shape [9].

However, it is based on linear measurements and has little use for reconstructing the full shape. In most cases, spore shape is treated as a qualitative trait. Spore shape terminology in mycology is a traditional system with dozens of terms that use common geometric shape categories (“globose”, “cylindric”) or similarity to some natural or cultural objects (“ovoid”, “filiform”). The picture becomes further complicated when subcategories are arbitrarily introduced, e.g. via adding prefix “sub-” or epithets “slightly” or “almost”. The subjectivity of this system represents a problem for reproducible research. Furthermore, such an approach makes impossible an assessment of the variation of a trait on an individual, populational, or species level. In their review, [1] concluded that “geometric morphometrics in the study of fungal shapes was far less employed compared to other microscopic organisms”.

To our knowledge, only two studies analysed the shape of fungal spores quantitively. Study [11] found differences in the spore shape between the populations of one species and study [12] – between the populations as well as different species. However, neither of these mycological studies nor most other morphometric studies questioned in a quantitative way how the object’s shape performs compared to size when identifying species. An answer to this question would help to decide how much effort should be invested into digitizing shape information for such small objects as fungal spores. In this study, we test the hypothesis that shape, as a richer source of information, overperforms size when performing automated identification of fungal species. If it is the case, fungal spore shapes are worth the digitizing effort and integration into taxonomic workflows. To test this hypothesis, we are going to answer the following questions:

i. Is it possible to adequately extract the shape information from images of fungal spores with the software that was developed for macroscopic objects?
ii. Can the shape differences found in multivariate analyses be reconstituted as the outlines and recognized by a human eye?
iii. Is it justified to split the spore shape information into symmetric and asymmetric portions or analysis of the overall shape variation would suffice?

As a test system for our study, we will use the genus *Subulicystidium* Parmasto (Hydnodontaceae, Trechisporales, Agaricomycetes, Basidiomycota, Fungi). It is a genus of fungi with 22 known species [13]. All known *Subulicystidium* species have smooth crust-shaped (corticioid) fruiting bodies and occur as saprotrophs on moderately or strongly decayed wood and are common in many forest ecosystems, especially tropical ones. For this genus, we own a large set of spore images after our previous study [14]. In that study, we performed traditional morphometric analysis of the spores while treating spore shapes only qualitatively. An additional advantage of our test system is the availability of DNA sequences for numerous specimens and the availability of the detailed genus-level phylogenetic tree. The DNA-based species assignments will serve as reference information to compare the performance of shape versus size data for automated species identification.

## Methods

### DNA data

We used a balanced set of 30 herbarium specimens which included ten species of *Subulicystidium* and where each species was equally represented by three specimens. We treated two clades of *S. perlongisporum* described in [15] as two separate species “*S. perlongisporum* 1” and “*S. perlongisporum* 2”. For all 30 specimens, we isolated and sequenced the internal transcribed spacer (ITS) of the nuclear ribosomal DNA as described in protocol 1 in [15] or used the sequences we produced earlier [14]. We processed raw sequence data with Geneious version 5.6.7 [16]. We imported the edited sequences to R version 4.0.3 [17] with the package “Biostrings” version 2.58.0 [18] and performed the multiple sequence alignment with the package “msa” version 1.18.0 [19] using MUSCLE algorithm [20] and other settings as default. We then customly trimmed the ends of the alignment with the package “ips” version 0.0.11 [21] to the length when 15 of 30 sequences had non-ambiguous base characters in the first and last position of the alignment. The resulting alignment had 734 nucleotide positions. We used the alignment viewer available in the package “ape” version 5.4.1 [22] to check visually the sequence alignment. According to the Akaike criterion [23], we identified “TrN+G+I” as the best-fitting nucleotide substitution model. We seacrhed for the best-scoring maximum likelihood phylogenetic tree with the nearest neighbor interchange strategy and performed bootstrap analysis (1000 replicates) with the “phangorn” package version 2.5.5 [24]. We visualized the result with the R package “ggtree” version 2.4.1 [25].

We submitted the newly generated DNA sequences to GenBank [26]. We provide the full list of the used DNA sequences with GenBank accessions and metadata on voucher specimens in supporting information S1.

### Morphological data

#### Spore terminology

To describe the spore morphology, we used the terms as they are found in [27] and [28]. The spores in *Subulicystidium*, as in all Basidiomycota, are produced externally on a sporangium called basidium and remain attached to it till they become mature and ready for discharge (Fig 1). It is the proximal part of the spore that directly contacts the basidium. The distal part of the spore is found on the opposite side of its long axis. The spore has an adaxial side, i.e. turned to the main axis of basidium, and opposite to it an abaxial side. On the proximal part of the spore, there is a projection called hilar appendix that is involved in the spore discharge from a basidium [27]. Observing hilar appendix on the adaxial side of the spore means the spore is seen in the lateral face. Observing hilar appendix directly on the main axis of the spore means the spore is seen in the frontal face. It is correct to compare the shapes of the spores within the same face. In our study, we focus on the spore’s lateral face which is more informative in the case of *Subulicystidium*.

**Fig 1.**
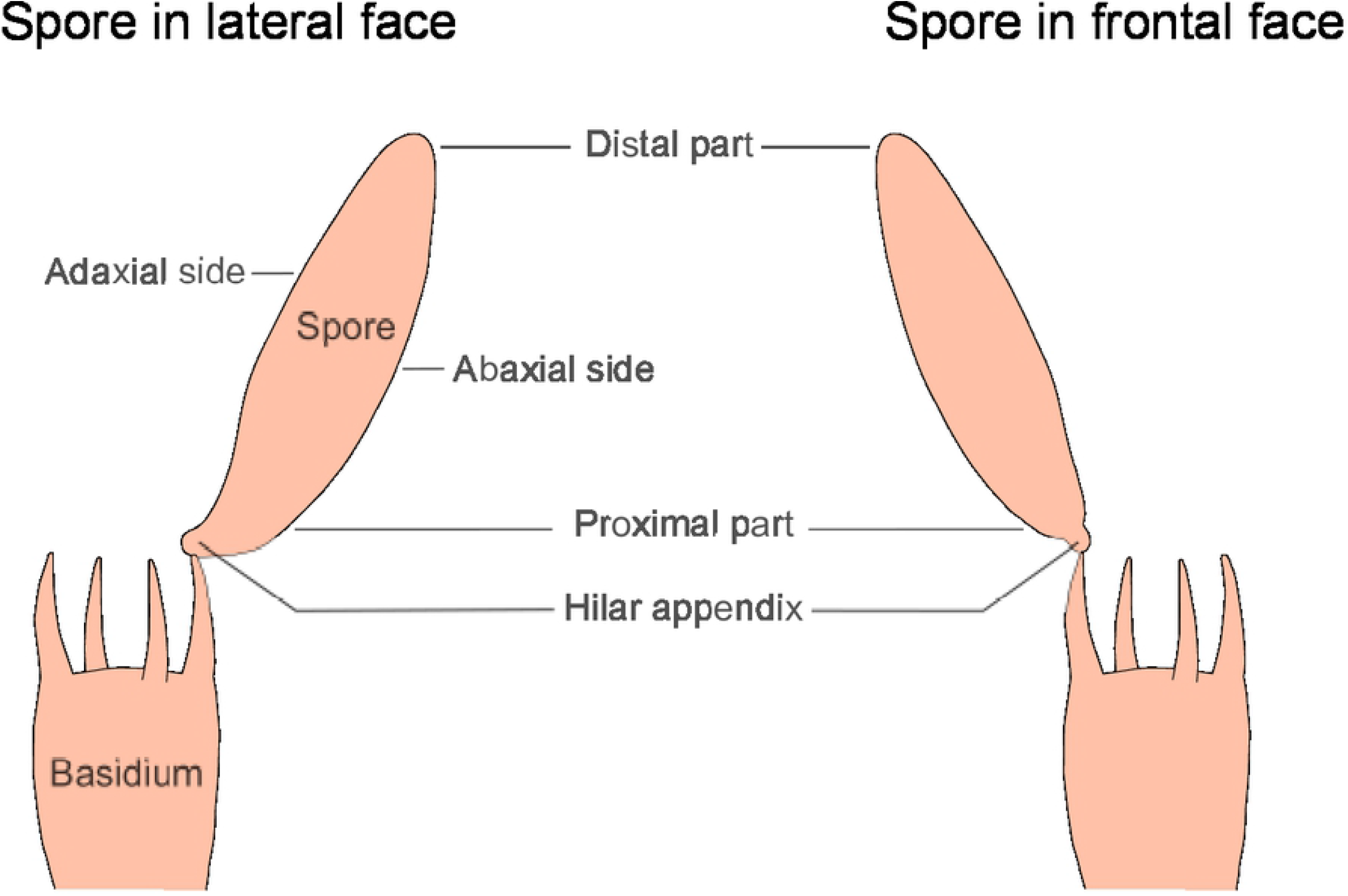
Crucial terms for describing a spore of the member of Agaricomycetes.

#### Image acquisition and pre-processing

We acquired and pre-processed images from light microscopy as described in detail in our online protocol [29]. In this paper, we highlight the most essential steps and we illustrate a workflow of image processing in Fig 2. We performed all work on images on a desktop computer with 64-bit Windows 10 operating system (build 19041). We obtained images of spores from squash preparations of fungal herbarium specimens examined at 1000× magnification (Fig 2A). The size of the captured images (JPEG files) was 1024 × 768 or 2048 ×1536 pixels while the resolution was always 96 dpi. We performed bulk image renaming with Bulk Rename Utility version 3.3.1.0 [30] and conversion from JPEG to BMP graphic format with ImageMagick version 7.0.10- [31].

**Fig 2.**
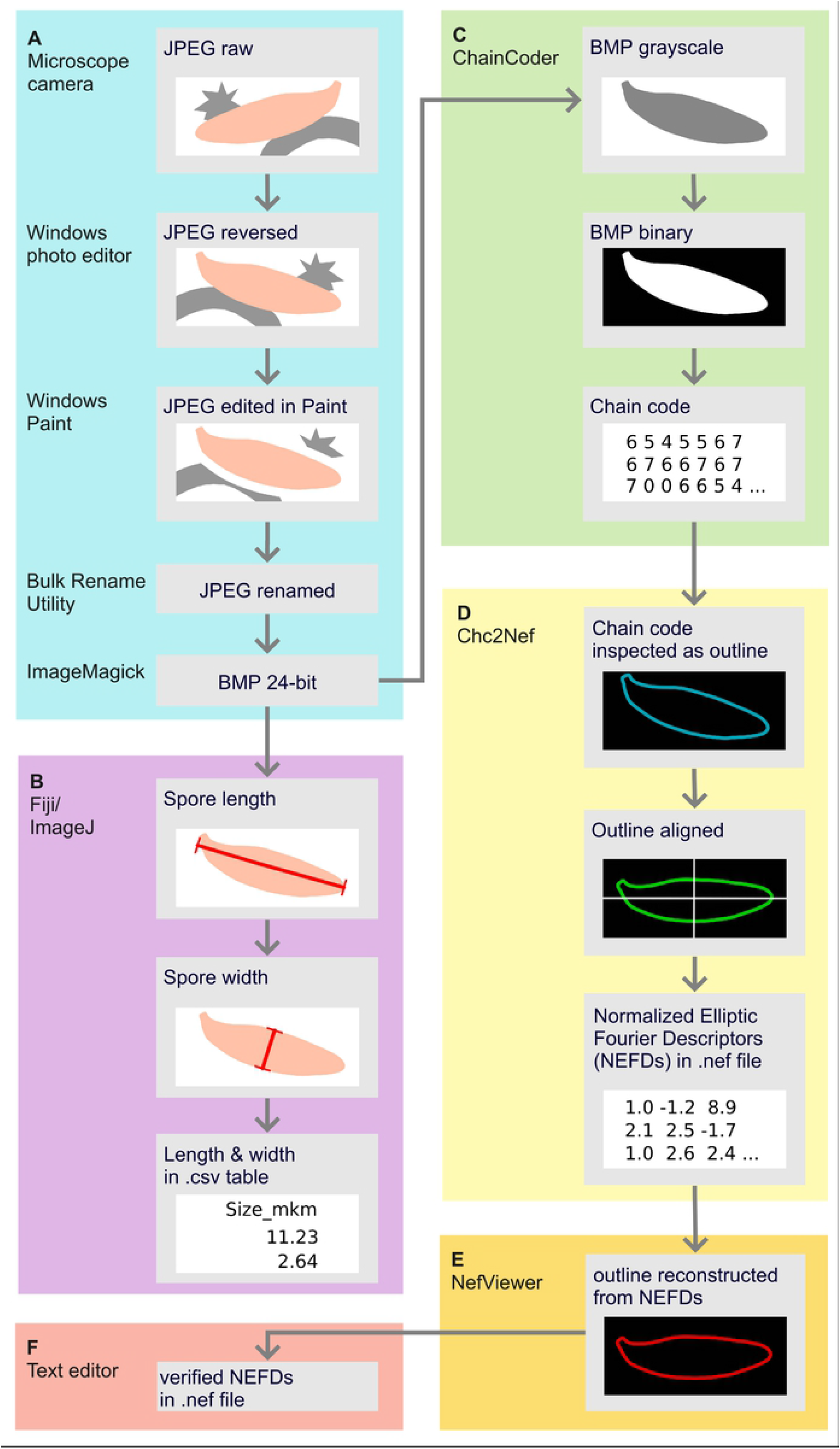
Workflow showing the extraction of the shape and size information from a fungal spore. (A) Image capture and pre-processing, (B) Acquiring linear measurements (C) Acquiring chain codes (D) Transforming chain codes to Normalized Elliptic Fourier Descriptors (E) Checking quality and orientation of the outlines (F) Possibility to open and manage outline descriptors in a text editor. The tools and programs used at each step are named on the left side of each colored panel.

For further processing, we selected those images that contained one or several healthy (not broken, with intact cell wall) mature (not attached to basidia) and well-focused spores [29]. We selected also only the spores that were laying clearly in the lateral face. Furthermore, for meaningful shape analysis, spores had to be alignable, i.e. with hilar appendix stretching out to the same direction if the spore were placed in the same orientation. Some images had to be mirrored to meet this criterion. In some cases, we applied additional manual adjustments, namely painting white lines around some parts of the spore outline, to enhance the clarity of spore outlines and to ensure the absence of contact with the structures that are other spores or not spores (crystals or hyphae), similar to [10] and [32]. These adjustments did not affect the geometric properties of the spore outlines but enabled a correct outline extraction. Raw and pre-processed images and all information extracted from them are available as a published dataset [33].

#### Processing shape information

Fungal spores are practically devoid of landmarks, i.e. distinct points that mark angles or cavities and can be unambiguously assigned to an object. Therefore, for fungal spores, methods aiming to reconstruct the complete outline are advantageous. Among such methods, Elliptic Fourier analysis [34] is most commonly employed. It uses an ellipse as a starting shape and searches for the coefficient values that transform the ellipse into the shape that reproduces the original outline of the object. These coefficients are called Elliptic Fourier Descriptors. After normalization, i.e. removing the effects of size, rotation, shift, and starting point of outline recording, they are called Normalized Elliptic Fourier Descriptors (NEFDs) and are used in statistical analysis as variables describing the shape [1].

We performed Elliptic Fourier analysis in the software package SHAPE version 1.3 [35] (http://lbm.ab.a.u-tokyo.ac.jp/~iwata/shape/). We used consequently each of the following programs from this package for different analysis steps: ChainCoder, Ch2Nef, NefViewer, PrinComp, and PrinPrint [29]. We started with grayscaling the BMP images in the program ChainCoder (Fig 2C). Then we converted the images to binary (assigning white pixels to spores and black to background) based on a threshold that was mostly automatically selected by ChainCoder and was only rarely adjusted manually. To remove the shape artifacts, we applied an erosion-dilation filter (that excludes the noisy pixels from the outline) and rarely also a dilation-erosion filter (to fill in artifact cavities). Then a chain code, i.e. a sequence of x and y coordinates describing the outline, was taken for each spore.

We imported the chain codes to Ch2Nef program [35] and checked the orientation of all spores to be the same, i.e. spore hilar appendix placed in the upper left quarter of the image (Fig 2D). When necessary, we applied 180-degree rotation to some of the spores. Ch2Nef then transformed the chain code into the Normalized Elliptic Fourier Descriptors (NEFDs). As a depth of detalisation of the contour information, we opted for 20 levels that are called harmonics in the elliptic Fourier analysis [34]. In general, other studies of organism shapes used from 15 to 30 harmonics [36,37], and using 20 harmonics was sufficient in the classical study from authors of the SHAPE package [6,35]. As each harmonic is described by four coefficients (= Elliptic Fourier Descriptors), there were 80 coefficients that characterised the shape of each spore. As the first three coefficients are always constant, there were 77 coefficients serving as shape variables for the next analysis step. To normalize the Elliptic Fourier Descriptors, we used the approach based on the first harmonic [35]. We checked that NEFDs are alignable with the NefViewer program (Fig 2E).

We combined manually NEFDs for individual specimens into a single .txt file (Fig 2F). To reduce the multidimensionality of NEFDs data, we run on it the Principal Component Analysis (PCA) in PrinComp program [35]. We used the flexibility of PrinComp and performed in total three principal component analyses (PCAs): i) considering only the symmetric shape variation, ii) considering only the asymmetric shape variation, and iii) considering overall shape variation not differentiated into symmetric and asymmetric (hereinafter called global shape variation). As a result of each PCA, we retained those principal components (PC) that explained a substantial amount of variation and were marked by PrinComp as having an eigenvalue >1. Such PCs were called “effective” in PrinComp. We inspected visually the shape variation accounted for by each effective PC with PrinPrint program [29]. The latter reconstructed the mean shape and plus and minus two times the standard deviation shape for each principal component [35].

#### Processing size information

For exactly those spores for which NEFDs were obtained, we took also the linear measurement: length (the longest dimension) and width (the broadest dimension) according to the standard accepted in mycology [9]. We used Fiji distribution of ImageJ version 1.53c [38] as shown in Fig 2B and described in [29]. Fiji does not differentiate whether the measurement is of a category “length” or “width”. Therefore, we measured constantly firstly length and then width or each spore. Though these values were originally placed in a single column, we split them into length and width columns in R using our protocol [29]. Based on length and width values, we also derived their ratio and used it as an additional variable in statistical analyses.

#### Comparative analyses

While obtaining the shape descriptors and linear measurements, we had control of the number of spores used per specimen and image. However, if the image contained several spores, we were not able to align the shape descriptors with linear measurements for each particular spore. Even though the input was the same .bmp image, the software for obtaining the shape data (SHAPE) and size data (Fiji) named individual measurements slightly differently. We could overcome it only by producing average-per-image trait values and using them as observations in correlations and discriminant analysis [29]. To produce average-per-image trait values, we used regular expressions in R as well as packages “stringr” version 1.4.0 [39] and “dplyr” version 1.3.0 [40].

For comparison, we assigned the morphometric traits to one of the five categories:

i. principal components of symmetric shape variation
ii. principal components of asymmetric shape variation
iii. principal components of global shape variation
iv. linear measurements (length and width)
v. length to width ratio

We explored how similar were all morphometric traits between themselves by Spearman correlations. To answer which morphometric traits allow more efficient automated identification of fungal species, we applied discriminant analysis. We assessed how accurate the individual traits and their combinations can identify (classify) to species level the observations whose species identity is not known to the statistical model. We found that trait values within species are not necessarily normally distributed and between the species do not have equal variance. For details, see supporting information S2 with the results of Shapiro-Wilk tests (made with R package “RVAideMemoire” version 0.9-79, [41]) and Levene tests (made with R package “heplots” version 1.3-8 [42]). Therefore, we applied an appropriate for such cases flexible discriminant analysis [43] implemented in “mda” R package version 05-2 [44]. We were randomly subsetting the data into the train (70%) and test (30%) portions attempting to balance the presence of data for different species with R package “caret” version 6.0-86 [45]. We repeated to subset the data into train and test portions 1000 times and then calculated the average identification (i.e. correct species assignment) success rate across subsettings using the R code adjusted from [46].

We managed data in R with the packages “here” version 1.0.1 [47], “conflicted” version 1.0.4 [48], “readr” version 1.4.0 [49], “data.table” version 1.13.4 [50], “dplyr” version 1.3.0 [40], and “report” version 0.2.0 [51]. We visualized the results with the packages „ggplot2” version 3.3.3 [52], “ggpubr” version 0.4.0 [53], function “cor.mtest” [54] and package “corrplot” version 0.84 [55]. We edited Figs 1 and 4 with Inkscape version 0.92.5 [56] and Fig 2 also with Miro online whiteboard (https://miro.com/). We provide the R project with the code, input data and results of analyses in supporting information S2 and as a repository on GitHub (https://github.com/ordynets/size_vs_shape).

## Results

Maximum likelihood phylogenetic analysis showed all ten *Subulicystidium* species on the phylogenetic tree as clearly separate clades, mostly with high bootstrap supports (Fig 3).

**Fig 3.**
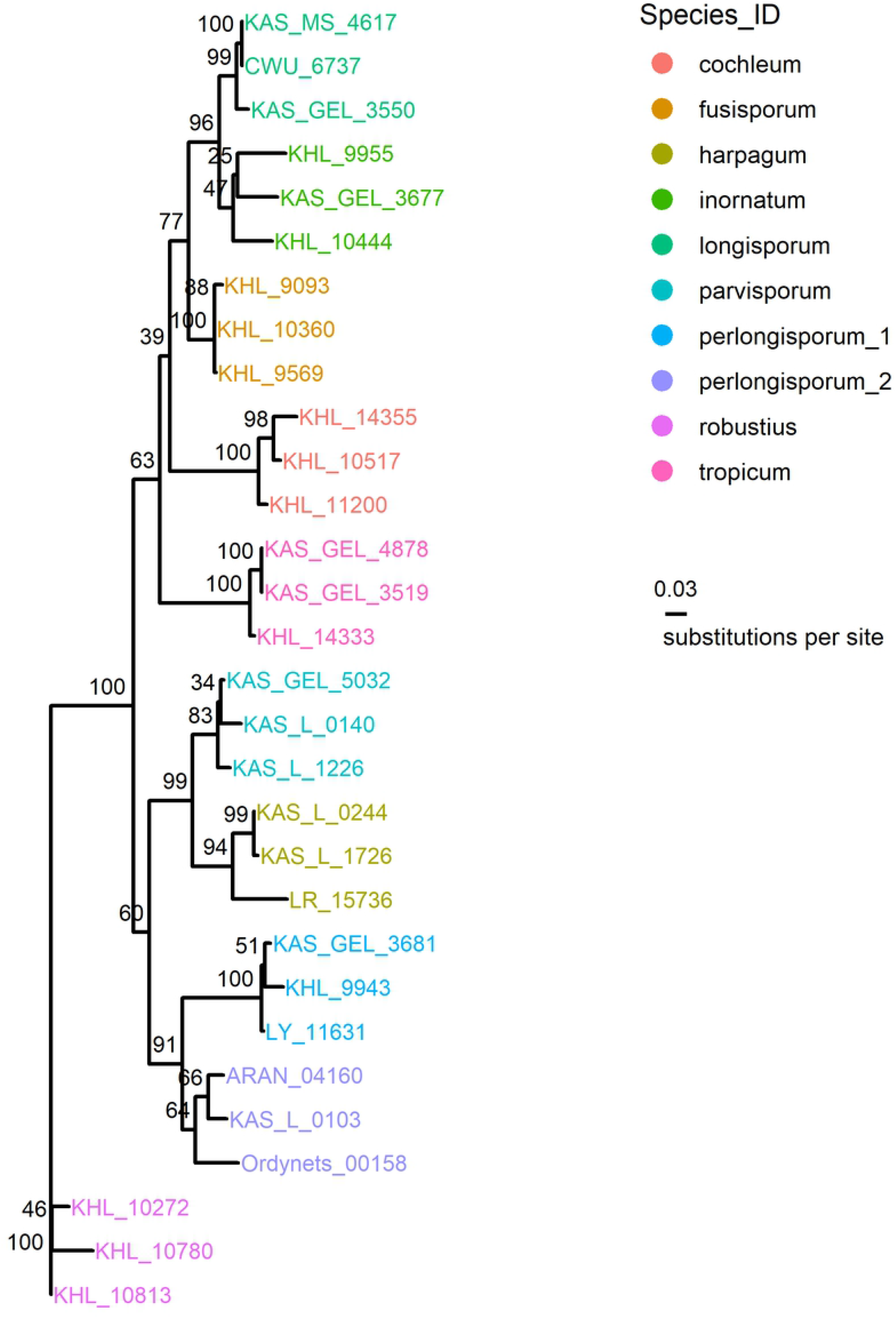
Phylogenetic relationship of 10 *Subulicystidium* species treated in this study.

After all quality filtering steps, we were able to analyse between 10 and 37 spores per specimen which totaled 511 spores from 30 specimens. These spores were captured in 401 images and each image contained one to four spores (supporting information S2). Therefore we considered 401 average-per-image values as observations for analyses.

PCA of symmetric shape variation identified only the first axis as effective (capturing 98,7% of the variation in data) while PCA of asymmetric variation three first axes (capturing 83.03, 6.63 and 4.11% of variation), and PCA of global variation two axes (capturing 94.57 and 2.91% of variation). Together with the spore length, spore width, and length to width ratio, this summed up to nine traits which we compared in terms of efficiency for automated species identification (supporting information S2).

By inspecting visually the shape variation accounted for by each principal component we found that the 1^st^ PC of symmetric variation reflected well the relative thickness of spores (Fig 4). The 1^st^ PC of asymmetric shape variation reflected the direction of single curving of the long axis of the spore. The 2^nd^ and 3^rd^ PCs of asymmetric shape variation reflected the variation in curvature of the proximal and distal end of the spore, respectively. The effects of the 1^st^ and 2^nd^ PCs of the global shape variation corresponded to the 1^st^ PC of symmetric and 1^st^ PC of asymmetric shape variation, respectively.

**Fig 4.**
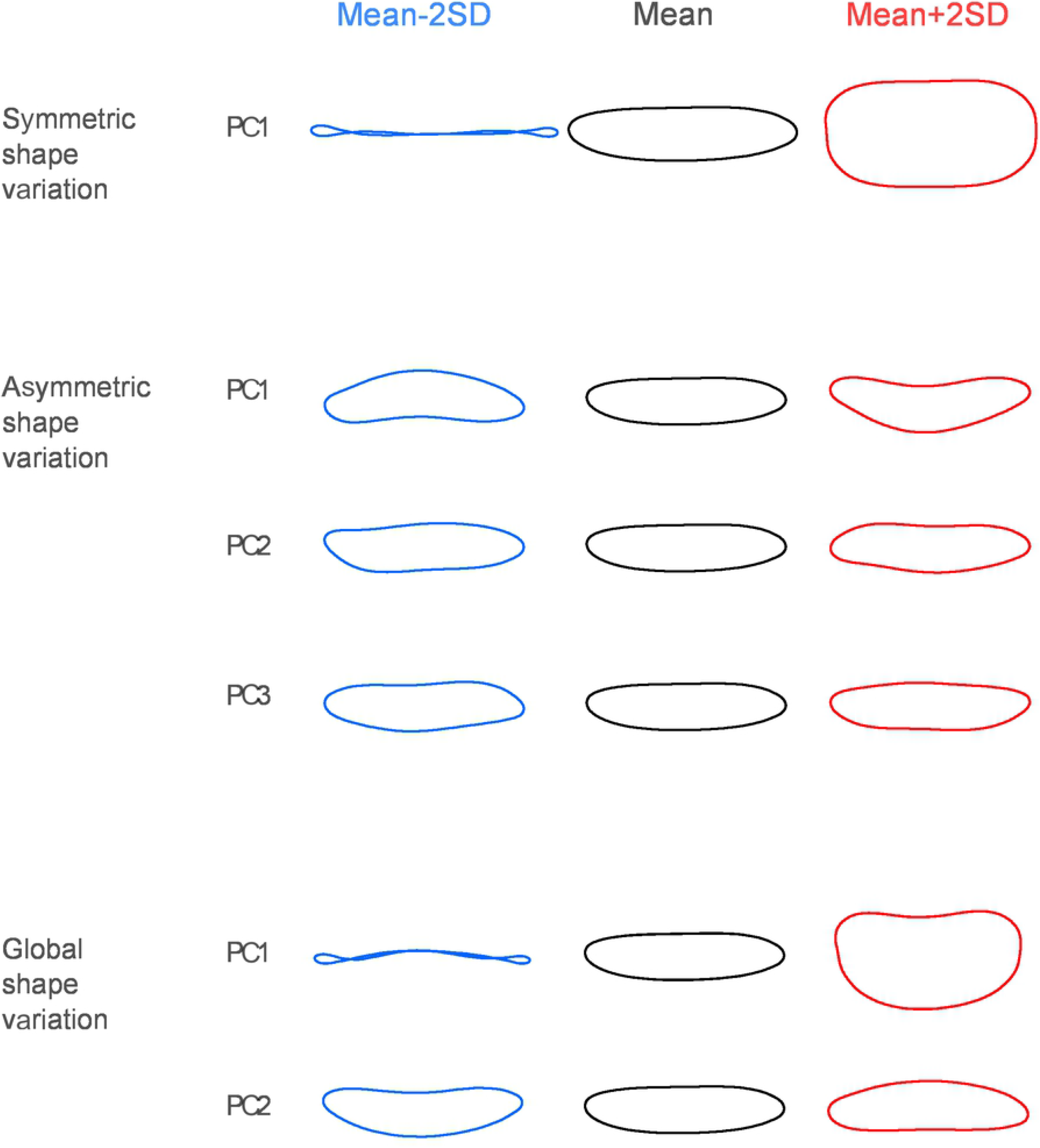
Ranges of shape variation that separate principal components account for.

When correlating the individual trait variables, we found that 1^st^ PCs of symmetric, asymmetric, and global shape variation correlated strongly positively with each other (Fig 5). All they correlated strongly negatively with the length and moderately positively with the width of the spores. The 2^nd^ PC of asymmetric shape variation correlated moderately negatively with the spore width while the 3^rd^ PC of asymmetric shape variation and 2^nd^ PC of the global shape variation did not correlate significantly with any other trait. Length and width correlated moderately negatively between themselves and each correlated very differently with the length to width ratio, though the correlation was stronger between the length and length to width ratio.

**Fig 5.**
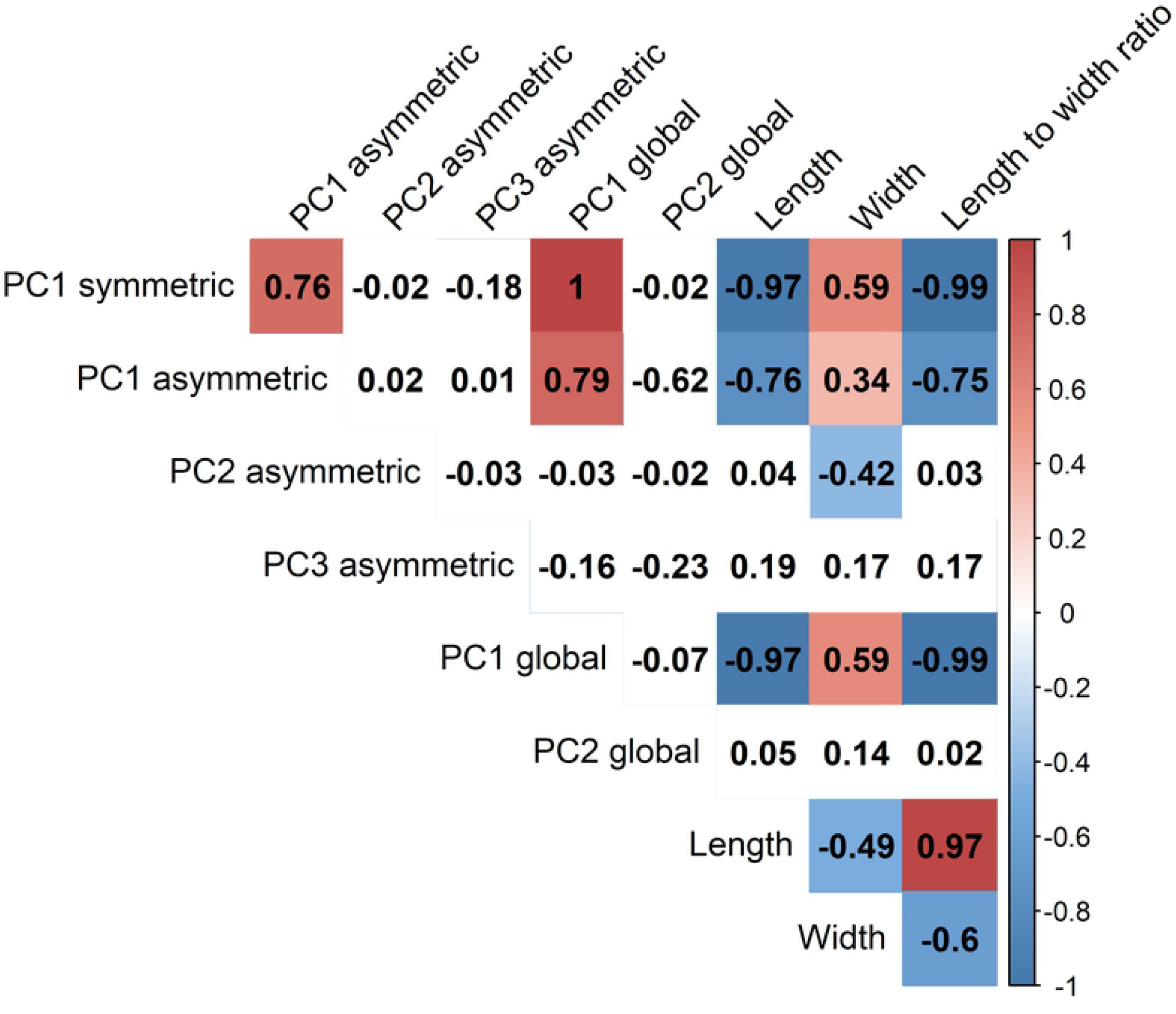
Spearman correlations of spore traits. Colors show the direction of correlation and highlight only significant correlation values (at level p=0.05)

In the discriminant analysis, the species identification success rate for the individual trait group was highest for the global shape variation (61.5%, Fig 6) while was slightly lower for length and width (59.1%). Symmetric shape variation identified the fungal species better than the length to width ratio (57.9% vs. 53.8%). The asymmetric shape variation had the lowest identification success rate, viz. 46.9%. When combining the traits of different groups, the highest identification success rate was achieved for the combination of symmetric, asymmetric shape variation and linear measurements (64.7%). By using the global shape variation instead of separate symmetric and asymmetric, the success rate was slightly lower (62.4%).

**Fig 6.**
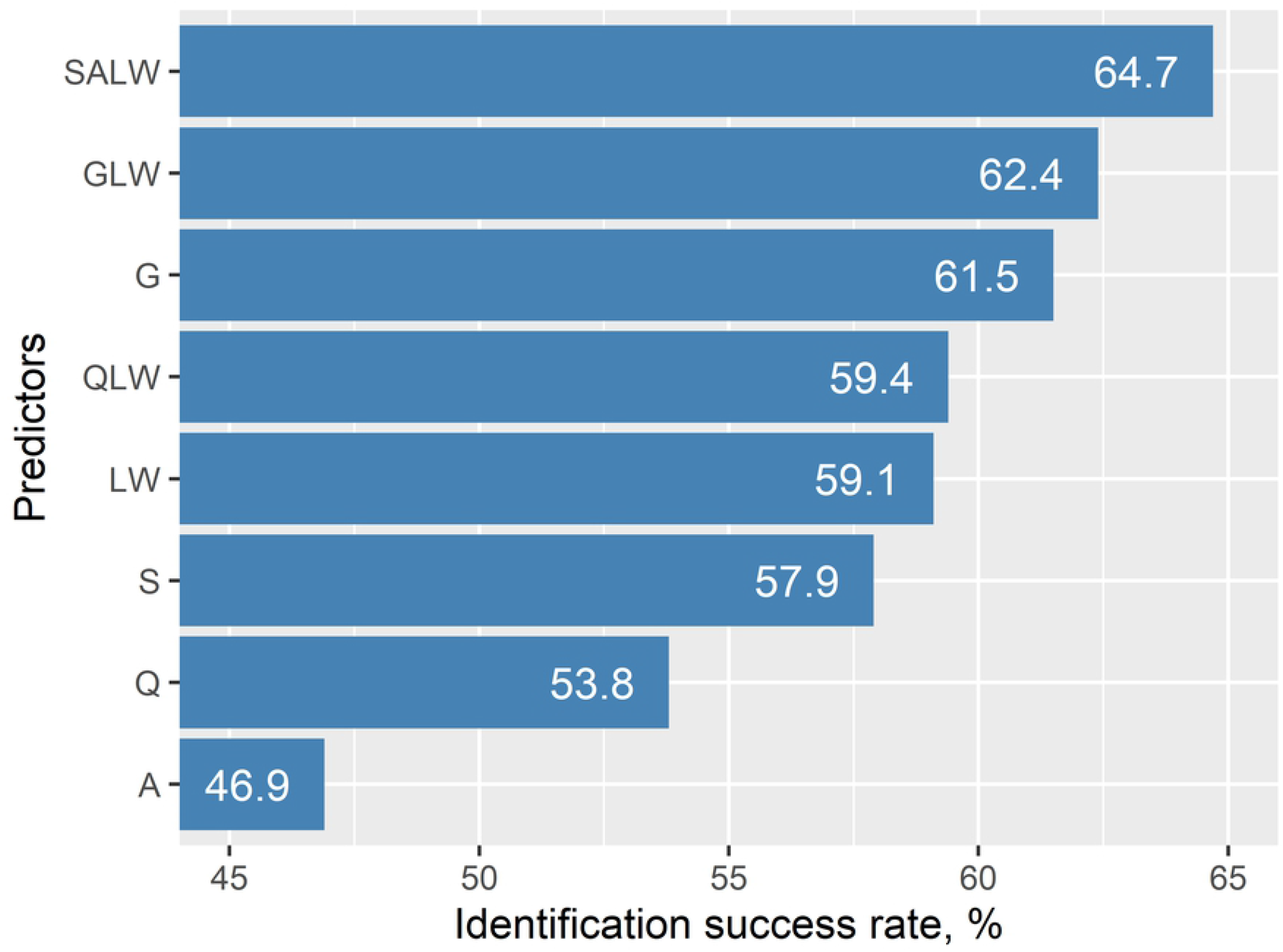
Species identification success rates in discriminant analysis for separate spore traits and their combinations. Abbreviations on y axis: S = symmetric shape variation (first principal component), A = asymmetric shape variation (three first principal components), G = global shape variation (two first principal components), LW = length + width, Q = length to width ratio, QLW = length to width ratio + length + width, GLW = global shape variation + length + width, SALW = symmetric shape variation + asymmetric shape variation + length + width.

## Discussion

Quantitative analysis of shapes helps to better identify and describe the organisms. In studies of fungi, despite their immense morphological diversity, quantitative shape analysis was almost ignored. In this study, we confirmed the hypothesis that shape, as a richer source of information, outperforms size during automated identification of fungal species by their spores. The highest identification success rate was achieved in a discriminant model that combined shape and size descriptors. The symmetric shape variation outperformed the classical length to width ratio. In general, we found that

i. It is possible to adequately extract the shape information from microscopy images of fungal spores;
ii. It is possible to recognise by a human eye the spore shape differences reconstituted after the multivariate analyses;
iii. It is justified to split the spore shape information into symmetric and asymmetric portions for separate analyses.

As quantitative shape analysis has been barely applied in mycology, we first had to ensure that available tools for extracting shape information can be used for our goal. Many of the tools were developed to work on the high-quality images of macroscopic organisms, often photographed as a single object per image in the desired orientation [57–59]. We opted for the program package SHAPE [35] which allows extracting several outlines from the same image, has flexibility for performing PCA and in general, has a convenient interface and detailed manual. Despite this package was originally designed for images of macroscopic objects, we successfully applied it to the outline extraction from the images with a modest resolution made by our microscope camera. The disadvantage of using SHAPE is a need to switch between the separate programs, meaning handling several outcomes and an extra effort in data management. Furthermore, the graphical user interface of SHAPE (or any other GUI program) means investing massive time effort when repeating the analysis and increased risk of producing an error. Therefore, in the future, it is advisory to work in a single environment with all analysis steps written as a code. This will simplify the data management and enhance the reproducibility of the analyses. Functions in R language designed by [60] and further developed by [61] and [62] are promising for this.

With our study design, we observed high correlations between several trait variables. The most important to note is the correlation between the shape descriptors (PC1 of symmetric and asymmetric variation) and size descriptors (especially length). This is due to our choice to represent the size as linear measurements, which are known to be geometrically dependent on a shape [2]. We kept linear measurements as size descriptors to demonstrate their properties and to explore the performance of these classical variables for species identification. We also generated the size variable that is a combination of separate linear measurements and is less independent of shape (square root of the product of length time width, as one possible option according to Claude 2008). This size variable indeed correlated less with shape variables but in discriminant model predicted the species much poorer than the length and width together (38.9% vs 59.1%, see supplementary information S2). In geometric morphometrics, there are also other measures of the size that are independent of shape and are worth the look by fungal taxonomists [60,63]. Among other trait variables, the correlation between PC1 of global and symmetric shape variation, PC1 of global and asymmetric shape variation, and length and length to width ratio are easily explained by their nested nature. Therefore we did not add these highly correlating variable pairs to the same discriminant models. On the other hand, we struggled to explain the rather high correlation between PC1 of symmetric and asymmetric shape variation. We guess that it is due to a strongly dominating character of PC1 of symmetric shape variation, which was constructed with an account of the first-level elliptic Fourier harmonic (D1).

The identification success rate in our discriminant analyses reached the maximum value of 64.69%. It is hard to compare our results with other studies where authors used different traits and/or different classification methods. Benyon et al. (1999) achieved an identification success rate of 66.2% using seven traits and a higher rate (74.4%) using 20 traits of fungal spores (mostly size descriptors). Authors of [4] obtained similar to our results in discriminant analyses of size and texture traits in diatom algae (55.1% correctly identified items). Authors of [65] reached 100% or nearly such rates for species of diatom algae by shape and texture descriptors. However, they used discriminant analysis as a dimensionality reduction tool and above its result applied other classification techniques. Authors of [66] evaluated their analyses of diatom cell shapes visually, i.e. as strength of separation of observations in the space of discriminant functions. They concluded that taxa with distinct shapes were separated well while the taxa with similar shapes were separated to a less extent. The identification success rates in [32] for statoliths in cubomedusae were comparable, after cross-validation, with ours, but were highly dependent on the number of observations per group. Due to technical reasons explained in Methods section, we performed analyses on average-per-image data. We admit that working with spore-level data would provide a more precise estimate of the correct identification rate.

Future work on quantitative shape analysis of fungal spores should cope with several challenges. Fungal spores may have very different outline properties in lateral versus frontal face. In the current study, we focused on the lateral face which is more peculiar in *Subulicystidium* and allows to capture the curvature along the spores’ long axis (while the frontal face masks this feature). However, in other fungal taxa, the lateral and frontal faces are both important [28] and their shapes should be analysed with the same attention. Authors of [32] combined three faces of the objects to achieve a high identification success rate. In fungi, it may be easy to identify the spore as exposed in the lateral face if its hilar appendix is large enough. Unfortunately, this is not the case for all species. Special care should be taken to identify the position of the hilar appendix in the needle-like spores (as in *S. cochleum* and *S. perlongisporum* in our dataset). Applying scanning electron microscopy may help to identify correctly the hilar appendix but also to get a detailed picture of the surface of the fungal spore. While our study focused on the fungi with smooth spores, the latter in many taxa bear additional projections like warts or ridges or distinct spines. These elements would be difficult to capture with the elliptic Fourier descriptors because of the mathematical properties of the method [1]. Therefore, ornamented spores of fungi give a chance to bring another approach, landmark-based methods of geometric morphometric, to mycology. These can be implemented in a two-dimensional and even three-dimensional space. Finally, different properties of the spores in lateral versus frontal face, as well as the availability of the several spore types per species in many fungal lineages, offer a possibility to bring the machine learning techniques to mycology.

We conclude that quantifying fungal spore shapes is worth the effort. We provide an open access protocol to propagate a broader use of quantitative shape analysis in fungal taxonomy and to stimulate the development of more efficient solutions to address the challenges we discussed above.

## Data Availability

Metadata on studied voucher specimens is provided in supporting information S1. DNA sequences newly generated for this study are available from GenBank as accessions <u>MW711723-MW711729</u>. Raw and pre-processed images and all information extracted from them are available as a published dataset (Ordynets et al. 2021b, https://doi.org/10.15156/BIO/807451). The R project with the code, input data, and results of analyses is provided in supporting information S2 and as a repository on GitHub (https://github.com/ordynets/size_vs_shape).

## Acknowledgements

We acknowledge the curators of fungal collections in ARAN, CWU, GB, LY, and O for providing specimens for our study, and personally Karl-Henrik Larsson, Janett Riebesehl, and Manuel Striegel who collected most of the specimens. David Scherf, Ludmila Lysenko, Ilka Kellner, Robert Liebisch, and Jonathan Denecke performed the DNA lab work and helped with the microscopy of the collections. Anton Savchenko, Oleh Prylutskyi, and Iryna Yatsiuk provided feedback on the earlier version of the protocol for extracting the shape and size information from the spores.

## Supporting information

S1 Table. GenBank accessions of used DNA sequences and metadata on voucher specimens

S2 Appendix. R project with the code, input data, and results of analyses.

